# Encoding of motor sequences in primate globus pallidus and motor cortex: Uniform preference for ordinal position

**DOI:** 10.1101/2025.09.15.676392

**Authors:** Devin R. Harsch, Karin M. Cox, Kevin K.M. Wright, Patrick J. Rice, Benjamin Pasquereau, Robert S. Turner

## Abstract

How the brain organizes discrete actions into fluid sequences is a central problem in motor neuroscience. Competing models of basal ganglia (BG) function propose that BG neurons either signal sequence boundaries or encode movements across ordinal positions. Prior studies have largely examined fixed sequences with end-of-sequence rewards, leaving open whether such findings generalize to more naturalistic conditions. We trained four rhesus macaques to perform a visuomotor sequence task requiring four or five out-and-back joystick movements to peripheral targets. Sequences were completed under two conditions: a random condition, in which target order varied across trials, and a fixed condition, in which order was predictable and consistent.

Rewards were delivered after each movement, dissociating reward timing from sequence completion. We recorded single-unit activity in arm-related regions of the globus pallidus (GP; n = 458) and primary motor cortex (M1; n = 306). Regression analyses revealed that many neurons in both GP and M1 encoded ordinal position within a sequence. Order effects were more frequent in the fixed condition, but were also present during random sequences. We found no evidence for preferential encoding of sequence initiation or termination in overlearned sequences, in contrast to prior studies reporting start/stop signals in basal ganglia. Weak effects appeared under the random condition in one animal pair, but these did not generalize across animals or conditions. Instead, neurons exhibited heterogeneous order-related responses spanning the full sequence. These results demonstrate that GP neurons, like those in M1, encode ordinal position throughout a sequence rather than acting solely as sequence initiators or terminators. This challenges boundary-specific models of BG function and highlights the BG’s broader role in representing serial order during motor sequence production.

## INTRODUCTION

How the brain chains together individual movements into fluid, efficient motor sequences is a central question in neuroscience known as “the serial order of motor behavior problem.” (Rosenbaum, Cohen et al. 2007) This ability to produce highly stereotyped sequences effortlessly is crucial not only for the expression of innate behaviors such as grooming (Berridge, Fentress et al. 1987) but also for the execution of highly practiced skills (Krakauer, Hadjiosif et al. 2019, Verwey 2024). How do the different components of cortical and sub-cortical motor control circuitry work together to achieve such feats? An influential series of studies in mice reported that neurons in the dorsolateral striatum selectively modulate their activity at the boundaries of a well-learned sequence of movements (Jin and Costa 2010, Jin, Tecuapetla et al. 2014), “bracketing” the start and the end of the sequence (Graybiel 1998). Those, and supporting results from studies in NHPs (Kimura, Kato et al. 1996, Hikosaka, Nakamura et al. 2002), are often cited as evidence for a model of basal ganglia function in which output from the BG serves to initiate and terminate well-learned motor programs. Yet, other studies have not observed start-or stop-related activity, finding instead that striatal neurons monitor ongoing kinematics of movement (Sales-Carbonell, Taouali et al. 2018, Dhawale, Wolff et al. 2021, Mizes, Lindsey et al. 2023). Studies in primates have found that individual neurons in the internal segment of the globus pallidus (GPi) selectively encode specific ordinal positions within a sequence of actions (Mushiake and Strick 1995), without showing a preference across the population for the start or stop of the sequence.

Neurons in the primary motor cortex (M1) have also been reported to show sequence-selective activity (Ben-Shaul, Drori et al. 2004, Lu and Ashe 2005, Matsuzaka, Picard et al. 2007). Those studies did not report how much of that was bracketing-type activity, as would be expected if bracketing activity was a common characteristic of the whole cortical-subcortical neural circuit for sequence production. Other studies, however, have failed to substantiate the presence of sequence-selective activity in M1 (Yokoi, Arbuckle et al. 2018, Zimnik and Churchland 2021). If sequence-selective activity and, more selectively, bracketing activity is concentrated preferentially within BG circuits, then that would suggest a unique role for BG circuits in sequence production.

The divergent results concerning the presence of sequence-selective and bracketing activities in the BG and M1 likely arise from differences between studies in task design, the amount of practice received by the subject, the specific subregion sampled, and analysis methods used. An additional potential source of confounds worth highlighting here is the possible influence of reward delivery. Although motor sequence learning and production in the real-world often occurs in the absence of explicit primary rewards (Verwey 2024) studies of sequential motor behavior in non-human subjects almost always motivate performance by the strategically-timed delivery of primary rewards – typically at the successful completion of the sequence.

Reward-related neuronal activity, especially that of dopaminergic neurons (Schultz 2017), has been found to encode reward prediction error in response to the delivery of primary rewards and also, critically, occurrence of the earliest predictor of reward delivery. (Additional influences of higher order reward contingencies and intrinsic goals have also been described (Gadagkar, Puzerey et al. 2016) but are beyond our scope here.) Thus, modulations of neuronal activity that appear to bracket the beginning and end of a motor sequence may instead be attributable to neural encoding of reward prediction error, especially in areas heavily innervated by dopaminergic neurons such as the BG. To gain a deeper understanding of the neural correlates of sequence production it is important to dissociate those from the potential encoding of reward prediction error.

We trained four nonhuman primates on two versions of a sequential reaching task (two animals each) to test the hypothesis that basal ganglia neurons selectively encode sequence boundaries, and to determine if that preference extends to neurons in M1. Animals moved a hand-held joystick through a series of out-and-back movements to capture a sequence of on-screen targets. In separate blocks of trials the targets appeared in random order or in an immutable order that the animals had practiced for months prior to data collection. Rewards were delivered upon completion of each sequence component thereby reducing potential confounding influences of reward prediction error. We sampled the spiking activity of single-units in the GP and M1 and used a general linear model to control for movement kinematics and to examine how those neurons encode ordinal position within random and learned fixed movement sequences.

We found no evidence for a preferential encoding of the beginning and/or end of motor sequences, but rather found a full gamut of the possible order-specific responses. These results suggest that preferential encoding of sequence boundaries is not a common feature in the GPi, primary output nucleus of BG skeletomotor circuit, nor in M1. Furthermore, although coding of ordinal position was more prominent during the production of well-practiced fixed sequences, it was also present on trials in which the target presentation order was randomized. We conclude that neurons in both GPi and M1 encode specific aspects of learned motor sequences thus suggesting roles for both of these structures in solving the serial order problem of motor behavior.

## MATERIALS AND METHODS

### Animals, Apparatus, and Tasks

Four monkeys (*Macaca mulatta*) were used for this study: one male (monkey H, 12 kg) and three females (monkey C, 7.5 kg; monkey F, 5.5 kg; monkey E, 6 kg). Procedures were approved by the Institutional Animal Care and Use Committee and complied with the Public Health Service Policy on the humane care and use of laboratory animals.

An animal was seated in a primate chair and held a joystick at the height of the mid-sternum (Fig. 1A). An LCD monitor was positioned vertically in front of the animal. Voltages reflecting the X and Y position of the joystick were digitized at 1 kHz. A 1-cm displacement of the joystick reflected a 1-cm displacement of a 3-mm white cursor on the screen, allowing the animal to control the on-screen position of the cursor. Empty gray circles (radius 20 mm) were shown on the screen: one center “start position” circle and either 4 or 8 potential peripheral targets. Four peripheral target zones (45°, 135°, 225°, 315°; hereafter referred to as targets 1–4) were used with monkeys C and H. Eight targets (0°, 45°, 90°, 135°, 180°, 225°, 270°, 315°; hereafter targets 1–8) were presented to monkeys E and F.

**Figure 1.**
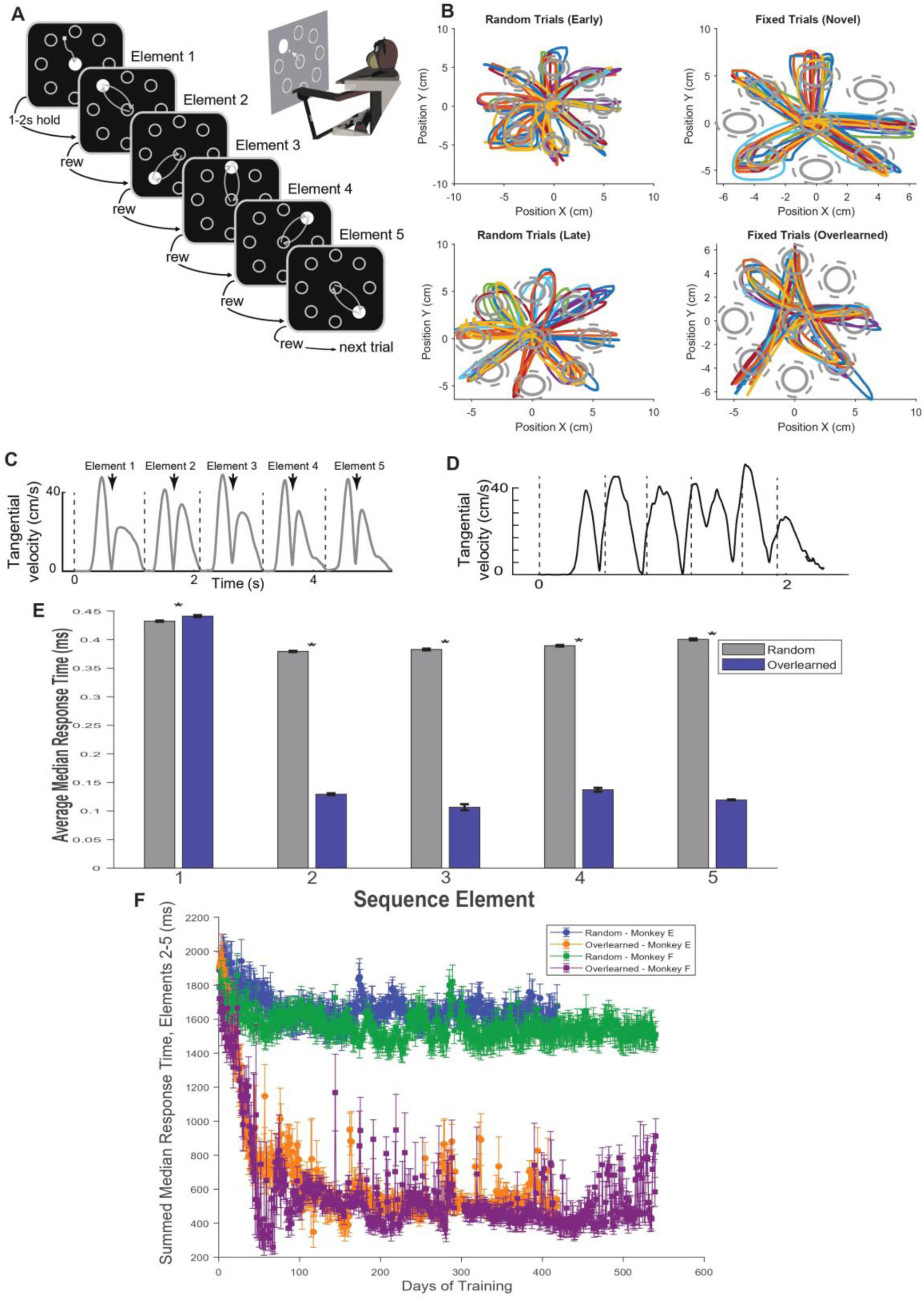
Animals learn to perform random and fixed motor sequences in a discrete sequence production (DSP) task. (A) **Task schematic.** Animals controlled a screen cursor with a joystick to perform a series of out-and-back reaches. A bolus of food reward was delivered after each successful reach. (B) **Cursor position traces.** Example trajectories from monkey E during random (left) and fixed (right) conditions, early vs. late in training. Each colored trace represents one successful trial. Top left: random trials from the first week of training. Bottom left: random trials after 400 days of training. Top right: fixed trials on the first day of fixed sequence exposure. Bottom right: fixed trials after 400 days of training. Trajectories become progressively more stereotyped with training. (C) **Velocity profile, random sequence.** Example tangential velocity trace from a single random-sequence trial in monkey E. Dashed lines mark returns to the central start position. (D) **Velocity profile, fixed sequence.** Same as in (C), but for a fixed-sequence trial. (E) **Response times by sequence element.** Average median response time across all recording days for random (gray) and fixed (blue) sequences, separated by sequence element. Asterisks indicate significant differences (p < 0.05). (F) **Learning curves.** Summed median response times for sequence elements 2–5, plotted across training days for monkeys E (random = blue, fixed = orange) and F (random = green, fixed = purple). Performance in the fixed condition improves markedly over training.

In this experiment, animals performed a variant of the discrete sequence production (DSP) (Verwey 2024) task adapted for visuomotor reaching (Fig. 1), which required multiple out-and-back movements between the central start position and either four (monkeys C and H) or five (monkeys E and F) peripheral targets. Many aspects of the paradigm have been described in detail in previous reports (Desmurget and Turner 2008, Desmurget and Turner 2010, Ramkumar, Acuna et al. 2016). A single trial progressed as follows (Fig. 1A): *1)* At the start of a trial, a filled white circle (“instruction cue”) appeared within the central start position, instructing the animal to move the cursor into this area and hold it there for a randomized time period (1-2 s, uniform random distribution). *2)* The filled white instruction cue appeared in one of the peripheral target zones, instructing the animal to initiate a movement toward this target. *3)* Immediately upon cursor entry into the peripheral target, the instruction cue appeared back in the central start position, prompting the animal to return the cursor to that area. *4)* When the cursor returned into the start position a drop of food reward was delivered to the animal via a sipper tube and the instruction cue appeared immediately in the next peripheral target. Stages 2-4 of the task were repeated four (monkeys C-H) or five (monkeys E-F) times in quick succession until the last out-and-back movement of the series was completed. When the animal’s task performance did not meet the timing or spatial requirements of the task, the trial was aborted and a 1-s inter-trial interval ensued without reward. The animals performed sequences under two conditions: a random condition, in which the target presentation order varied from trial to trial, and a fixed condition, in which the order was identical across trials (hereafter referred to as the “overlearned” condition). For monkeys C and H, the Overlearned target order was 1 → 2 → 3 → 4 and 4 → 2 → 1 → 3 (Monkeys C and H each performed two different Overlearned sequences, with data collection nonoverlapping between performance of the sequences). For monkeys E and F, the target presentation order was 6 → 3 → 8 → 1 → 4. All four animals initially practiced the Random version of the task in which the instructed peripheral target in step 2 was chosen at random with replacement. The animals performed the random out-and-back task a minimum of 6 months (>25,000 trials) prior to collection of the results presented here.

### Surgery

After training, each monkey was surgically prepared for intracranial recording by aseptic surgery under Isoflurane inhalation anesthesia. A cylindrical titanium recording chamber (18-mm ID) was affixed to the skull with stereotaxic guidance over a craniotomy contralateral to the arm used for the task. This allowed for transdural access to the arm-related regions of M1 and GP. The chamber was oriented parallel to the coronal plane at an angle of ∼35° so that electrode penetrations were orthogonal to the cortical surface. The chamber was fixed to the skull with bone screws and dental acrylic. Bolts were embedded in the acrylic to allow fixation of the head during recording sessions.

### Data acquisition

During the performance of the behavioral task, a glass-coated platinum/iridium (FHC, Bowdoin, ME; monkeys C-H) or tungsten (Alpha Omega, Nazareth, Israel; monkeys E-F) microelectrode was advanced into the site of interest (i.e., M1 or GP) using a hydraulic manipulator (MO-95, Narishige, Tokyo, Japan). Neuronal signals were amplified with a gain of 10K and bandpass filtered (0.3-10 kHz) and acquired either at 20 kHz (MAP, Plexon, Dallas, TX, USA; monkeys C-H) or at 25-kHz (RZ2, Tucker-Davis Technologies, Alachua FL; monkeys E-F). Individual spikes were discriminated on-line using Plexon off-line sorting software (Plexon Inc., Dallas TX). The timing of detected spikes and of relevant task events was sampled digitally at 1 kHz.

### Localization of the recording sites

To validate the anatomical location of the sites of interest (i.e., arm-related territories in M1 and GP) and proper positioning of the recording chamber, two methods were used: (1) Structural MRI scans (Siemens 3 T Allegra Scanner, voxel size of 0.6 mm) were performed for monkeys E-F. An interactive 3D software system (Cicerone) was used to visualize MRI images, define the target location, and predict trajectories for microelectrode penetrations. (Miocinovic, Zhang et al. 2007, Pasquereau and Turner 2013) (2) Alternatively, histologic reconstructions were performed after completion of recordings in monkeys C-H. Each animal was given a lethal dose of sodium pentobarbital and was perfused trans-cardially with saline followed by 10% formalin in phosphate buffer and then sucrose. Brain sections (50-µm) were processed histologically with cresyl violet staining to localize microelectrode tracks. By aligning microelectrode mapping results (penetrations spaced 1-mm apart) with MRI images or histologic reconstruction, we were able to estimate the anatomical location of microelectrode penetrations. The boundaries of brain structures were identified based on standard criteria, which included relative location, neuronal spike shape, firing pattern, and responsiveness to active or passive movements of contralateral joints (Turner and Anderson 1997). In addition, a cortical region was targeted for data collection if microstimulation at low currents evoked contraction of contralateral forelimb muscles (<40-µA, 10 biphasic pulses at 300-Hz).

### Analysis of behavioral data

X-and Y-position signals derived from the joystick were numerically filtered and combined to obtain kinematic measures such as hand position, velocity, speed, and acceleration. The time of movement onset/offset was defined as the point at which tangential velocity crossed an empirically-derived threshold (0.35 cm/s). Reaction times (RT: interval between ‘instruction cue’ and movement onset), movement durations (MD: interval between movement onset and capture of the target), peak speed (maximum speed), and movement extent (distance between movement onset and offset) were computed for each trial. All the data analyses were performed using custom scripts in the MATLAB environment (Mathworks).

### Analysis of Neuronal Data

Neuronal recordings were accepted for analysis based on electrode location, recording quality, and recording duration (>10 trials per peripheral target). Adequate single unit isolation was verified by testing for appropriate refractory periods (>1.5-ms) and waveform isolation (signal/noise ratio superior to 3 S.D.). The baseline firing rate for each unit was defined as the average spike count observed in the second prior to the presentation of the first peripheral target. For each unit, we counted spikes in a 200-millisecond window centered at the time of the peak velocity of the outward movement to a target. Thus, each successful trial yielded 5 spike counts – one for each reach to a peripheral target. Units with outlier firing rates were removed according to the following protocol: For each unit, firing rates were defined by pooling over the baseline and movement data. Firing rate outlier thresholds were determined with each of the three brain regions (GPi/GPe/M1) separately. For each region, we pooled across all monkeys (E/F/C/H) and identified inclusive upper and lower bounds, defining the maximum and minimum firing rates we would accept for that that region. These bounds were found by taking the pooled monkey data, and running the Matlab isoutlier function, using the default “median” method and a ThresholdFactor of 1.5. This procedure identifies the values corresponding to the median +/-1.5 scaled median absolute deviations. For each region, all units with FRs > the determined upper bound or < the lower bound were discarded. Following outlier removal, we applied a multiple linear regression analysis to investigate each retained unit’s response to key task features.

Specifically, we used the model:

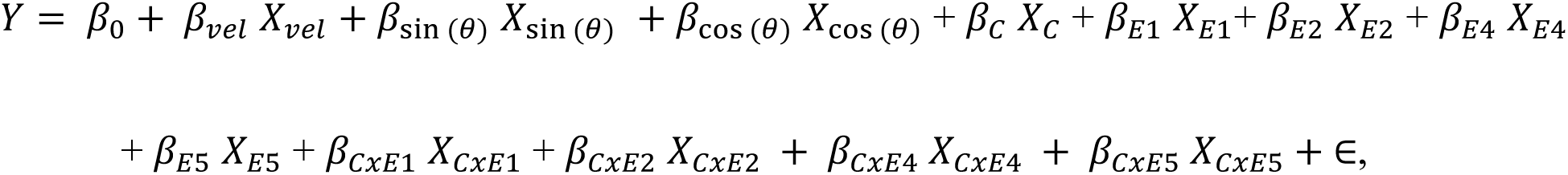

Where 𝑌 is the baseline-subtracted spike count, 𝛽_0_ is the intercept term, 𝛽_𝑣𝑒𝑙_ 𝑋_𝑣𝑒𝑙_is the peak velocity, 𝛽_sin (𝜃)_ 𝑋_sin (𝜃)_ + 𝛽_cos (𝜃)_ 𝑋_cos (𝜃)_ are the sine and cosine of the movement direction, 𝛽_𝐶_ 𝑋_𝐶_ is the main effect of condition, 𝛽_𝐸1_ 𝑋_𝐸1_+ 𝛽_𝐸2_ 𝑋_𝐸2_+ 𝛽_𝐸4_ 𝑋_𝐸4_+ 𝛽_𝐸5_ 𝑋_𝐸5_ represents the main effect of order, 𝛽_𝐶𝑥𝐸1_ 𝑋_𝐶𝑥𝐸1_ + 𝛽_𝐶𝑥𝐸2_ 𝑋_𝐶𝑥𝐸2_ + 𝛽_𝐶𝑥𝐸4_ 𝑋_𝐶𝑥𝐸4_ + 𝛽_𝐶𝑥𝐸5_ 𝑋_𝐶𝑥𝐸5_ represents the order x condition interaction, and ∈ is the normally distributed error. Note that for monkeys C and H, the model only goes up to E4, as they performed a four-target version of the task.

This analysis tested whether neuronal spike firing during motor sequence performance was influenced by kinematic parameters (e.g. movement direction and velocity, with assumed cosine tuning (Kennedy and Schwartz 2019), sequence order, task condition, or a condition x order interaction. We used indicator coding, with element 3 chosen as the reference level. Those units that exhibited a main effect of order or an interaction effect were then selected for subsequent analysis. We then used a Cohen’s f^2^ model to obtain the effect size for each unit for each condition (Cohen 1988). Units with an f^2^ statistic below 0.15 were excluded from further analysis. The variance associated with kinematic parameters was partialled out to obtain residual spike counts. The residuals were separated by task condition, and a subsequent linear regression was applied to assess condition-specific effects of order.

## RESULTS

### Performance of a fixed sequence becomes stereotyped after extensive practice

Animals were trained to perform the DSP task as described above (Figure 1A). Over the course of training, animals achieved highly skilled performance on the task. Figure 1B illustrates the difference in movement trajectories early versus late in training for the overlearned sequence. Each trajectory shows the position trace from a randomly selected correct trial. Late in learning, position traces become more stereotyped and efficient, consistent with expert-level performance (Ramkumar, Acuna et al. 2016, Ohbayashi 2021). This increase in efficiency is also evident in single-trial velocity profiles. Figure 1C shows the tangential velocity from a representative trial in the random condition for Monkey E. Each dashed line marks the animal’s return to the center target. In the random condition, animals return to the center target after completing the movement and pause until the next target appears. In contrast, this latency is absent in the overlearned condition (Figure 1D): animals return to the center and initiate the next reach without delay, often before the target is cued.

We calculated the average median response times for each cohort of animals for each sequence element and found significant differences across all elements. For monkeys E and F, the average median response time for element 1 was surprisingly higher for the first element (432.37 ms vs 441.13 ms, Random vs Overlearned, p <.001). For all other elements, the average median response time for the overlearned sequence was significantly lower (Element 2: 379.37 ms vs 129.25 ms, p <.001; Element 3: 382.77 ms vs 106.53 ms, p <.001; Element 4: 389.35 ms vs 137.07 ms, p <.001; Element 5; 400.50 ms vs 119.32 ms, p <.001; paired t-tests). Monkeys C and H showed a similar pattern for Element 2-4. Median response times were slightly shorter in the overlearned condition for element 1 (454.03 ms vs 427.33 ms, p <.001), and significantly shorter for elements 2–4 (Element 2: 428.67 ms vs. 252.32 ms, p <.001; Element 3: 443.95 ms vs. 184.19 ms, p <.001; Element 4: 463.42 ms vs. 293.47 ms, p <.001, paired t-test). To visualize the time savings over the course of the sequence, we summed the found the median response times for elements 2-5 of the sequence for each day of training (Figure 1F). Analogous plots to those shown in Figure 1 for Monkeys C and H can be found in Supplementary Figure 1. Together, these results confirm that animals exhibit expert-level performance on the DSP task.

### Single unit activity was recorded in primary motor cortex and the globus pallidus

We recorded neuronal activity of 458 GP (287 GPe, 171 GPi) cells from four animals, and 306 M1 cells from three animals. Recording sites for all four animals are available in Figure 2. All cells displayed some level of modulation during task performance. The results from these recordings are summarized in Table 1. Spike density functions for an example GP (Figure 3A and 3B) and M1 (Figure 3C and 3D) unit are shown in Figure 3.

**Figure 2:**
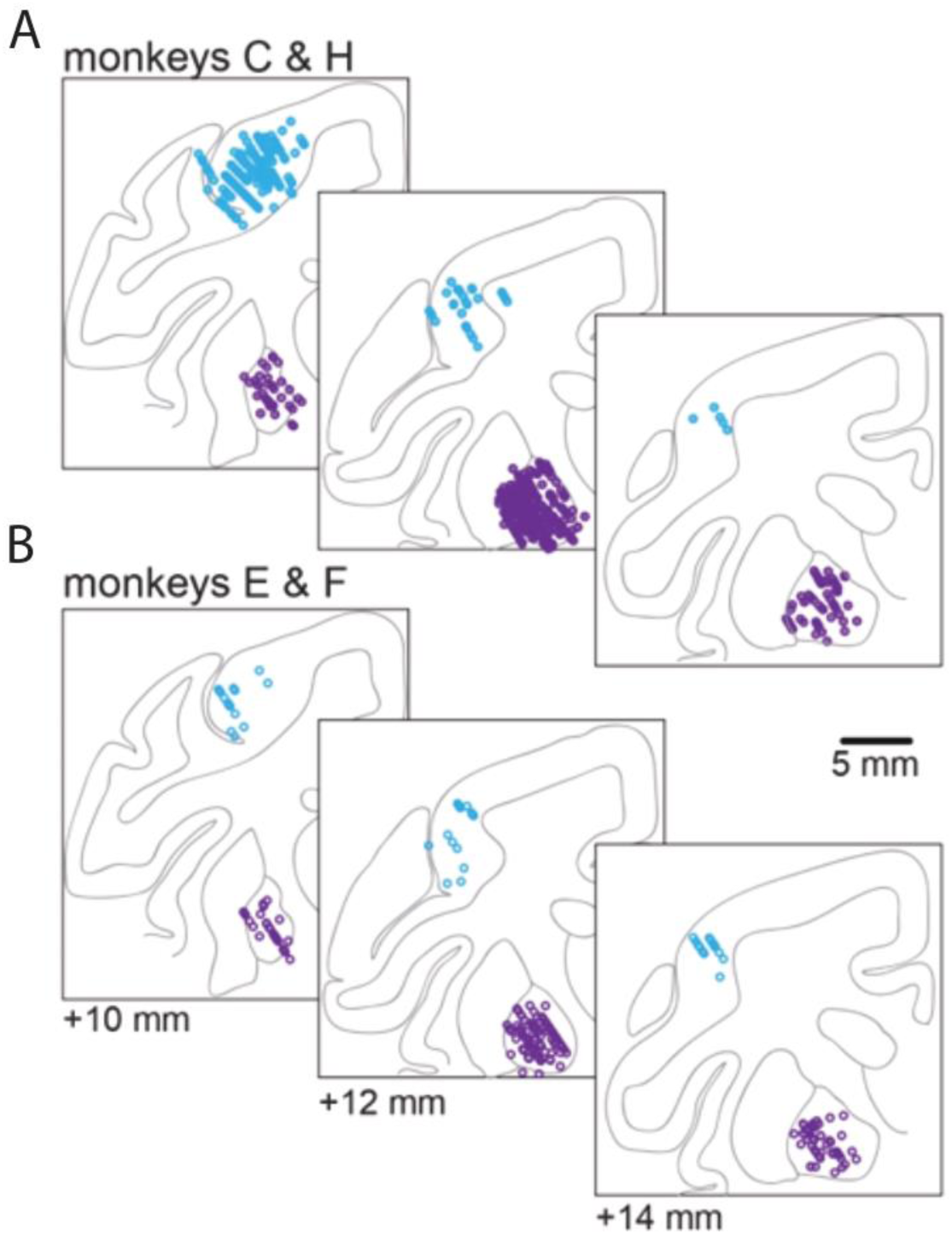
Reconstruction of recording sites. Blue dots indicate electrode penetrations in primary motor cortex (M1), and purple dots indicate penetrations in the globus pallidus (GP). (A) Recording sites for monkeys C and H. (B) Recording sites for monkeys E and F. Together these reconstructions confirm coverage of both cortical and basal ganglia regions across animals.

**Figure 3.**
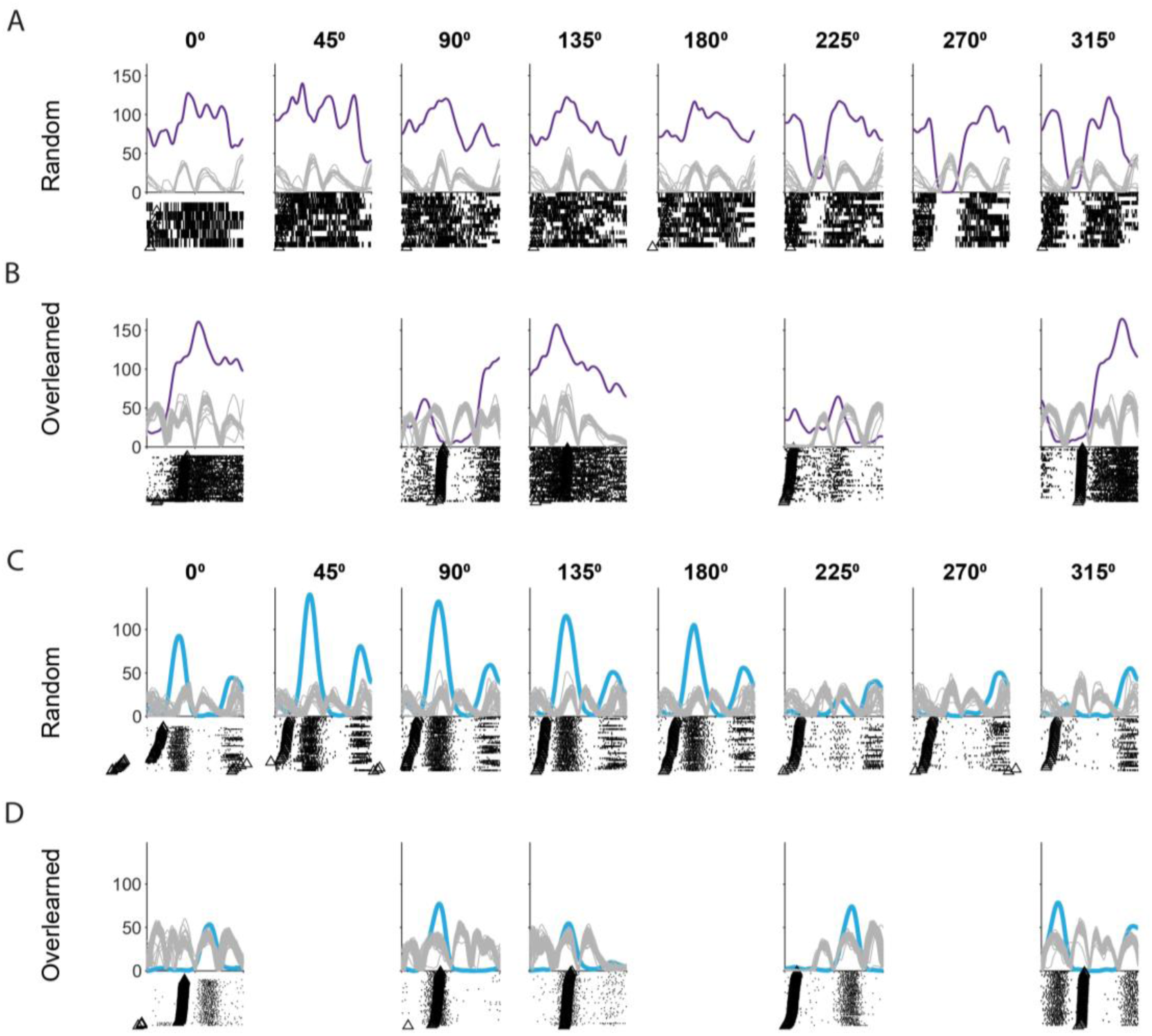
Example spike density functions (SDFs) and raster plots from GPi and M1 units during random and overlearned sequences. (A) GPi unit from monkey E in the random condition. Spike density functions (purple) are aligned to the end of the outward movement. Gray traces indicate tangential velocity profiles from individual trials, and black triangles mark the measured reaction time for each trial. Corresponding spike rasters are shown below, separated by target direction (0–315°). (B) Same unit as in (A), but during the overlearned condition. (C) M1 unit from monkey F in the random condition, plotted as in (A). Spike density functions (blue) are aligned to the end of the outward movement, with gray velocity traces and black triangles indicating reaction times.(D) Same unit as in (C), but during the overlearned condition.

**Figure 4:**
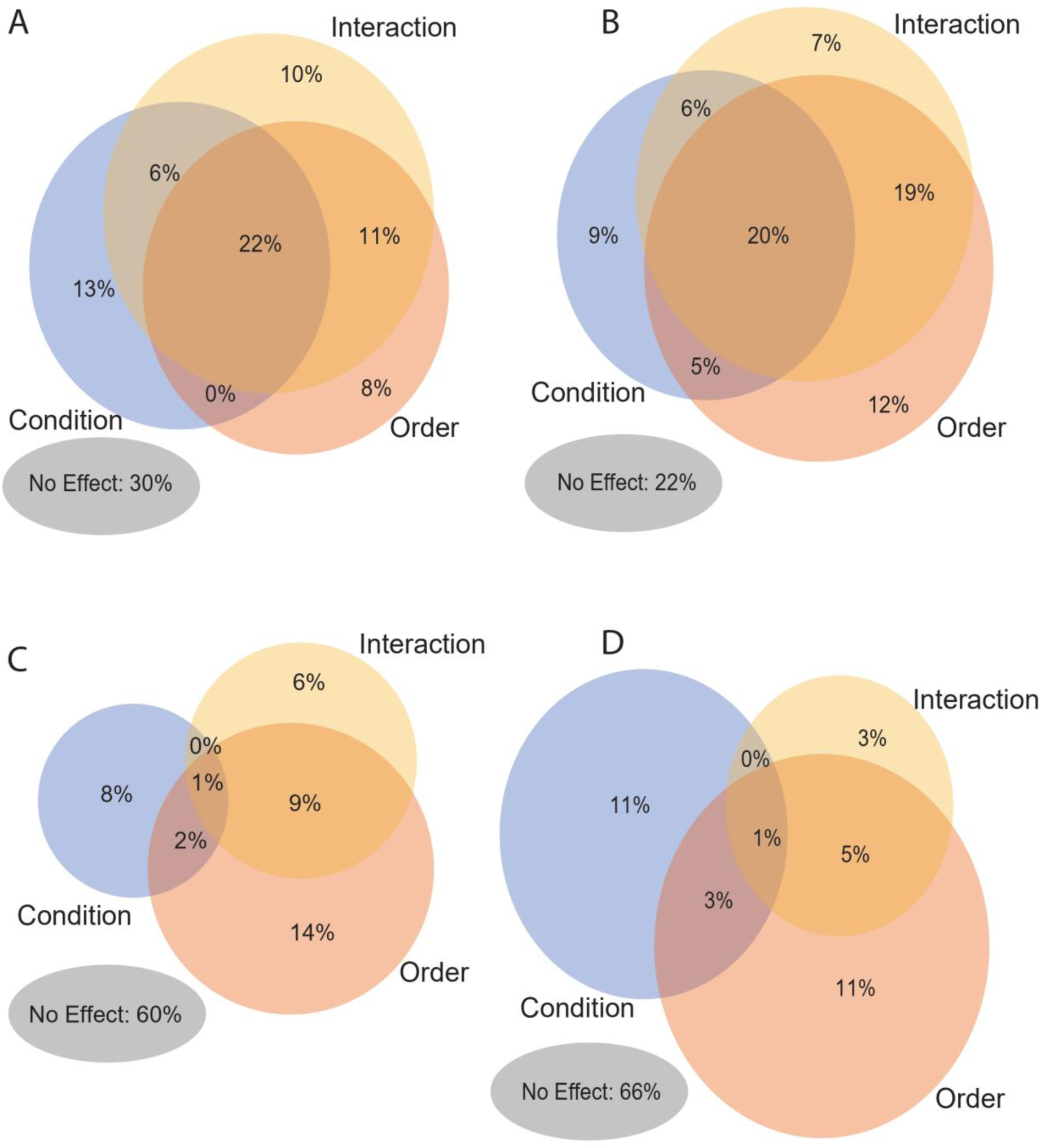
Venn diagrams indicating main effects as derived from general linear regression Each circle represents the proportion of units with a significant main effect of condition, order, or condition × order interaction. Overlaps indicate units with multiple significant effects. Percentages denote the proportion of the total recorded population in each category. (A) M1 units from monkeys E and F. (B) GP units from monkeys E and F. (C) M1 units from monkeys C and H. (D) GP units from monkeys C and H.

**Table 1:** Unit counts and average firing rates for recorded neurons. Values are mean ± SD. GPe: external segment of the globus pallidus; GPi: internal segment of the globus pallidus; M1: primary motor cortex.

| Monkey | GPe |  | GPi |  | M1 |  |
| --- | --- | --- | --- | --- | --- | --- |
|  | Unit count | Firing rate (Hz) | Unit count | Firing rate (Hz) | Unit count | Firing rate (Hz) |
| **E** | 37 | 81.19 ± 26.06 | 37 | 77.96 ± 25.90 | Not recorded | Not recorded |
| **F** | 45 | 92.86 ± 26.50 | 19 | 91.99 ± 28.54 | 72 | 11.55 ± 7.95 |
| **C** | 79 | 73.54 ± 25.66 | 37 | 81.38 ± 20.66 | 99 | 13.75 ± 8.94 |
| **H** | 126 | 89.09 ± 22.86 | 78 | 89.48 ± 22.18 | 135 | 12.41 ± 8.10 |

### A linear model dissociated neural encoding of kinematics and sequence

We counted the number of spikes in a 200 ms window centered at the peak velocity of the outward reach for each movement in the sequence. Linear regression was performed on baseline-subtracted versions values to determine main effects of order, condition, or an order x condition interaction, as well as the effects of movement direction and velocity. Units were selected for subsequent analysis if they displayed a main effect of order and/or an order x condition interaction. For monkeys E and F, linear regression revealed that 56% of GP units displayed a main effect of order, 41% of units displayed a main effect of condition, and 52% of units displayed an order x condition interaction (Figure 3B). For M1, 42% of units displayed a main effect of order, 40% of units displayed a main effect of condition, and 49% of units displayed an order x condition interaction (Figure 3A). Overall, 69% of GP units and 57% of M1 units were selected for subsequent analysis. For monkeys C and H, 20% of GP units displayed a main effect of order, 15% of units displayed a main effect of condition, and 9% of units displayed an order x condition interaction (Figure 3D). For M1, 26% of units displayed a main effect of order, 12% of units displayed a main effect of condition, and 17% of units displayed an order x condition interaction (Figure 3C). 23% of GP units and 33% of M1 units were selected for subsequent analysis.

To further understand the neuronal activity associated only with specific order effects, we partialed out the effect of movement kinematics to obtain residual spike counts. We then performed another regression separately for each condition on the residual spike counts to examine the condition-specific effect of order. To ensure that the effect size was significant, we calculated the Cohen’s f-squared value for each unit under each condition and selected for further analysis only those units with an f-squared value above.15. In monkeys E and F, 9% of GP units displayed a condition-specific effect of order under the random condition, and 62% of units exhibited this effect under the overlearned condition. In motor cortex, 3% of units displayed a condition-specific effect of order under the random condition, and 57% of units displayed this effect under the overlearned condition. In monkeys C and H, 9% of GP units displayed a condition-specific effect of order under the random condition, and 21% of units exhibited this effect under the overlearned condition. In motor cortex, 9% of units displayed a condition-specific effect of order under the random condition, and 31% of units displayed this effect under the random condition.

### Neuronal responses tile the sequence space in both GP and M1

We determined each unit’s preferred sequence element by identifying the element with the largest absolute residual spike count. To assess whether preferences were uniformly distributed or biased toward specific sequence positions (e.g., boundaries), we applied chi-squared goodness-of-fit tests (MATLAB).

In Monkeys E and F, GP units under the random condition exhibited a significant preference for the **first element** (p = 0.0315). However, under the learned condition, element preferences were uniformly distributed (p = 0.0738). Among Monkeys C and H, 9% of GP units showed a condition-specific effect of order, but these did not prefer any particular sequence position (p = 0.1826). Similarly, in the overlearned condition, 21% of GP units displayed a condition-specific order effect, with element preferences again uniformly distributed (p = 0.3022).

In Monkeys E and F, only 2 M1 units exhibited a condition-specific effect of order in the random condition after accounting for effect size (Cohen’s f^2^ > 0/15). 57% of neurons exhibited a condition-specific effect of order in the overlearned condition, encoding sequence elements uniformly (p = 0.185). Among Monkeys C and H, 9% of M1 units exhibited a condition-specific order effect with no element preference (p = 0.165). Under the overlearned condition, 31% of M1 units displayed a condition-specific effect of order, and this subset did not encode sequence elements uniformly (p = 0.001), showing an apparent preference for the third element of the sequence. These results contrast with prior findings in striatum and GPe, where neurons tended to modulate their activity primarily at the boundaries of motor sequences, with little modulation during internal elements (Jin and Costa 2010).

Strikingly, there was little difference in the proportions of units with a condition-specific effect of order between GP and M1, save for a significant difference in monkeys C and H in the overlearned condition (EF Rand: p = 0.0760, Chi-sqare = 3.1475; EF Overlearned: p = 0.4496, Chi-square = 0.5718; CH Rand: p = 0.9268, Chi-square = 0.0084; CH Overlearned: p = 0.0079, Chi-square = 7.0478).

For each unit, we normalized the vector of residual spike counts by dividing each element by the magnitude of the vector, making each vector into a unit vector with a magnitude of 1. We plotted these results by sequence element (Figures 5 and 6).

**Figure 5.**
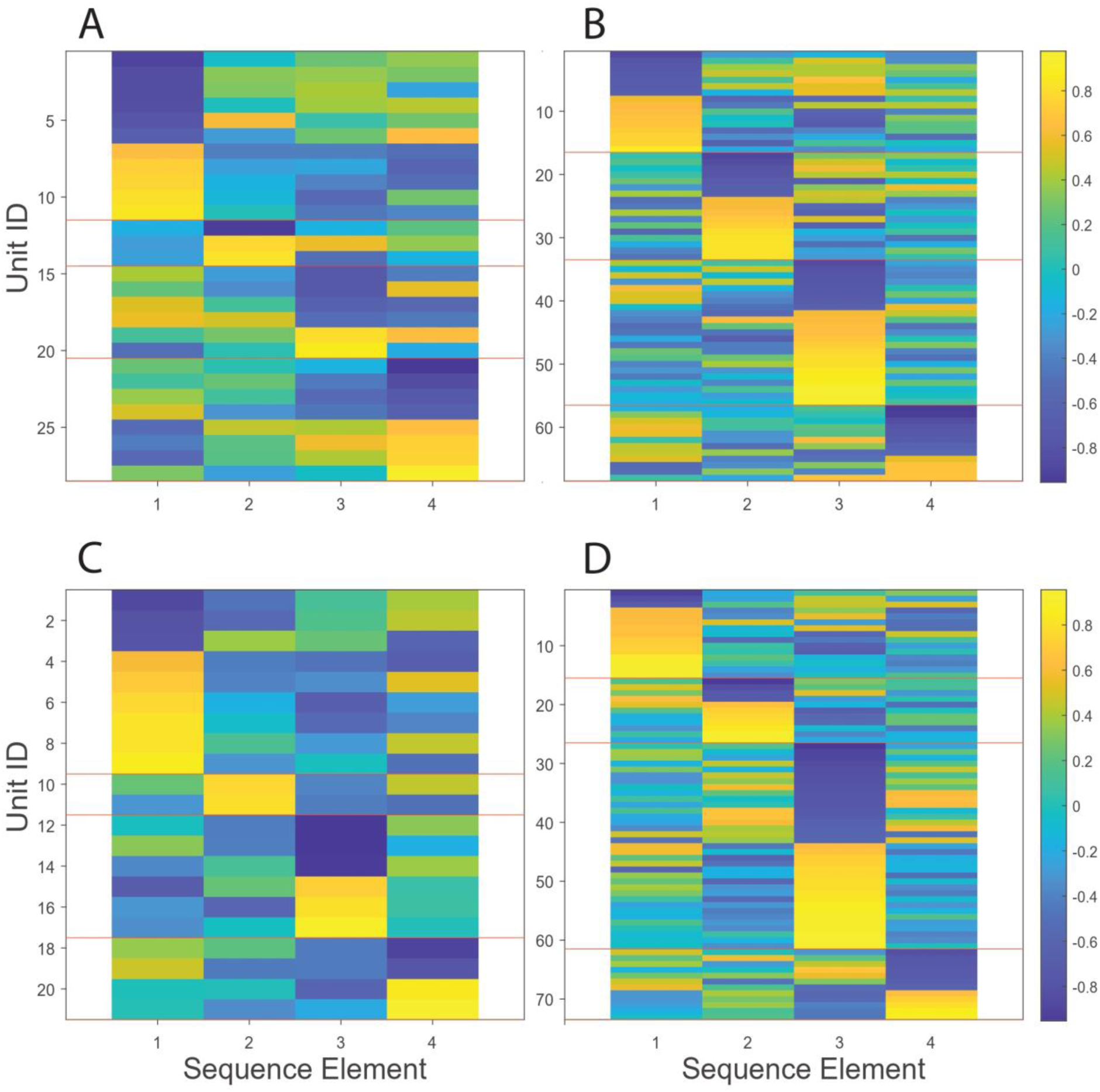
GP and M1 neurons in monkeys C and H exhibit heterogeneous order coding across sequence elements. Each row represents one unit and columns correspond to sequence elements (1–4). Color scale indicates normalized residual firing magnitude, with warm colors (yellow) denoting “increasing-type” responses (positive residuals) and cool colors (blue) denoting “decreasing-type” responses (negative residuals). Red horizontal lines indicate transitions between preferred sequence elements. Units are sorted by the sequence element of maximal modulation, revealing heterogeneous tiling of order selectivity across the population. (A) GP units, random condition. (B) GP units, overlearned condition. (C) M1 units, random condition. (D) M1 units, overlearned condition.

**Figure 6.**
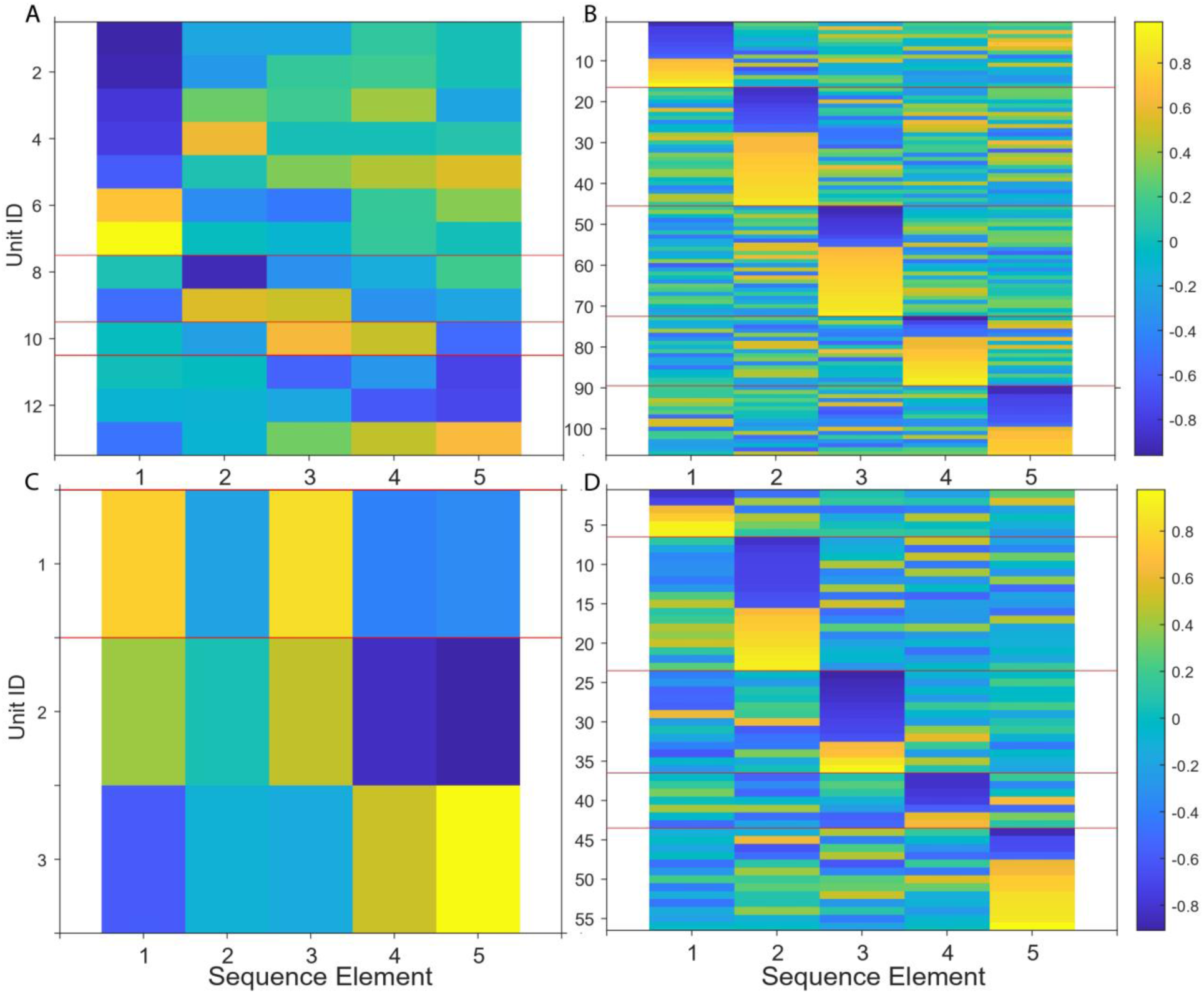
GP and M1 neurons in monkeys E and F exhibit heterogeneous order coding across sequence elements. Each row represents one unit and columns correspond to sequence elements (1–5). Color scale indicates normalized residual firing magnitude, with warm colors (yellow) denoting “increasing-type” responses (positive residuals) and cool colors (blue) denoting “decreasing-type” responses (negative residuals). Red horizontal lines indicate transitions between preferred sequence elements. Units are sorted by the sequence element of maximal modulation, revealing heterogeneous tiling of order selectivity across the population. Note that these animals performed a five-element sequence. (A) GP units, random condition. (B) GP units, overlearned condition. (C) M1 units, random condition. Only three units passed the Cohen’s f² threshold for effect size, with selectivity limited to sequence elements 3 and 5. (D) M1 units, overlearned condition.

### Response magnitudes are larger for the learned sequence

We next examined the magnitude of the neuronal responses by comparing the peak residual value for each unit and found that the response size was often greater for the learned condition than in the random condition. This is true for units that exhibit a condition-specific effect of order only in the learned condition, as well as for units that exhibit condition-specific order effects in both conditions. Furthermore, this heightened magnitude was present in both GP and M1 neurons. Given that we regressed out the effects of kinematics, this larger modulation is likely not due to differences in the way the animal performed the random and overlearned sequences. Instead, it suggests that neuronal responses in both GP and M1 are sensitive to task context. This modulation of activity in different contexts has been reported before in both GP (Mushiake and Strick 1995, Gdowski, Miller et al. 2001, Turner and Anderson 2005, Gdowski, Miller et al. 2007) and M1 (Omlor, Wahl et al. 2019, Mizes, Lindsey et al. 2024).

For GP neurons in monkeys E and F, 75% of units displayed a larger order effect for units with effects under both conditions, and 97% of units with an effect of order only under the learned condition displayed a larger order effect (Figure 6A and 6B). For M1 neurons in monkey F, 98% of units displayed a larger order effect for units with effects only under the learned condition (Figure 6C). Note that only three neurons exhibited an effect of order under both the learned and random conditions. We observed similar results in monkeys C and H (GP, both conditions: 85%; GP, effect in learned only: 99%; M1, both conditions: 83%; M1, effect in learned only: 99%; Figure 6d-g).

### Response patterns do not form separable clusters

Although the tiling effect we observed suggests that neurons in GPi and M1 encode sequence elements in a heterogeneous manner, it remained possible that these responses could be grouped into discrete categories defined by their activity across the sequence. To test this, we first performed principal component analysis (PCA) on the residual spike counts to examine whether units formed separable clusters in low-dimensional space. For both pairs of animals, PCA revealed no evidence of clustering in either GP or M1, under either the random or overlearned sequence conditions (Figure 7A–D). Instead, units were distributed smoothly throughout the space, consistent with a continuum of response types. Units recorded in the overlearned condition tended to show somewhat broader dispersion, but this reflected greater heterogeneity rather than the emergence of distinct groups.

**Figure 7.**
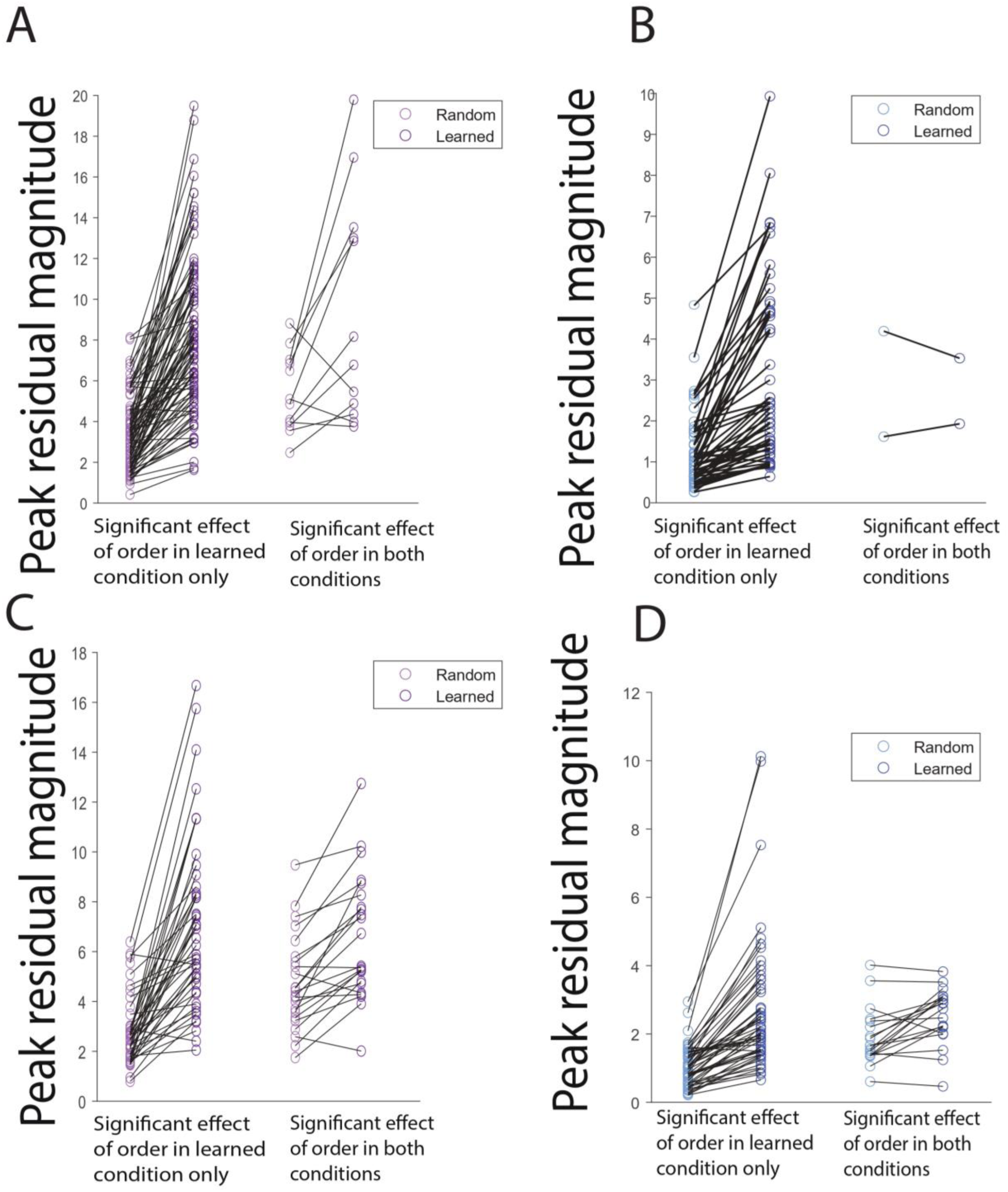
Response magnitudes are greater in the overlearned condition than in the random condition. Peak residual magnitudes for GP neurons in monkeys E and F, comparing random (gray) and overlearned (purple) conditions. Left: units with a significant effect of order in the overlearned condition only. Right: units with a significant effect of order in both conditions. (A) Same as (A), but for M1 neurons in monkeys E and F (C) Same as (A), but for GP neurons in monkeys C and H. (D) Same as (B), but for M1 neurons in monkeys C and H

**Figure 8.**
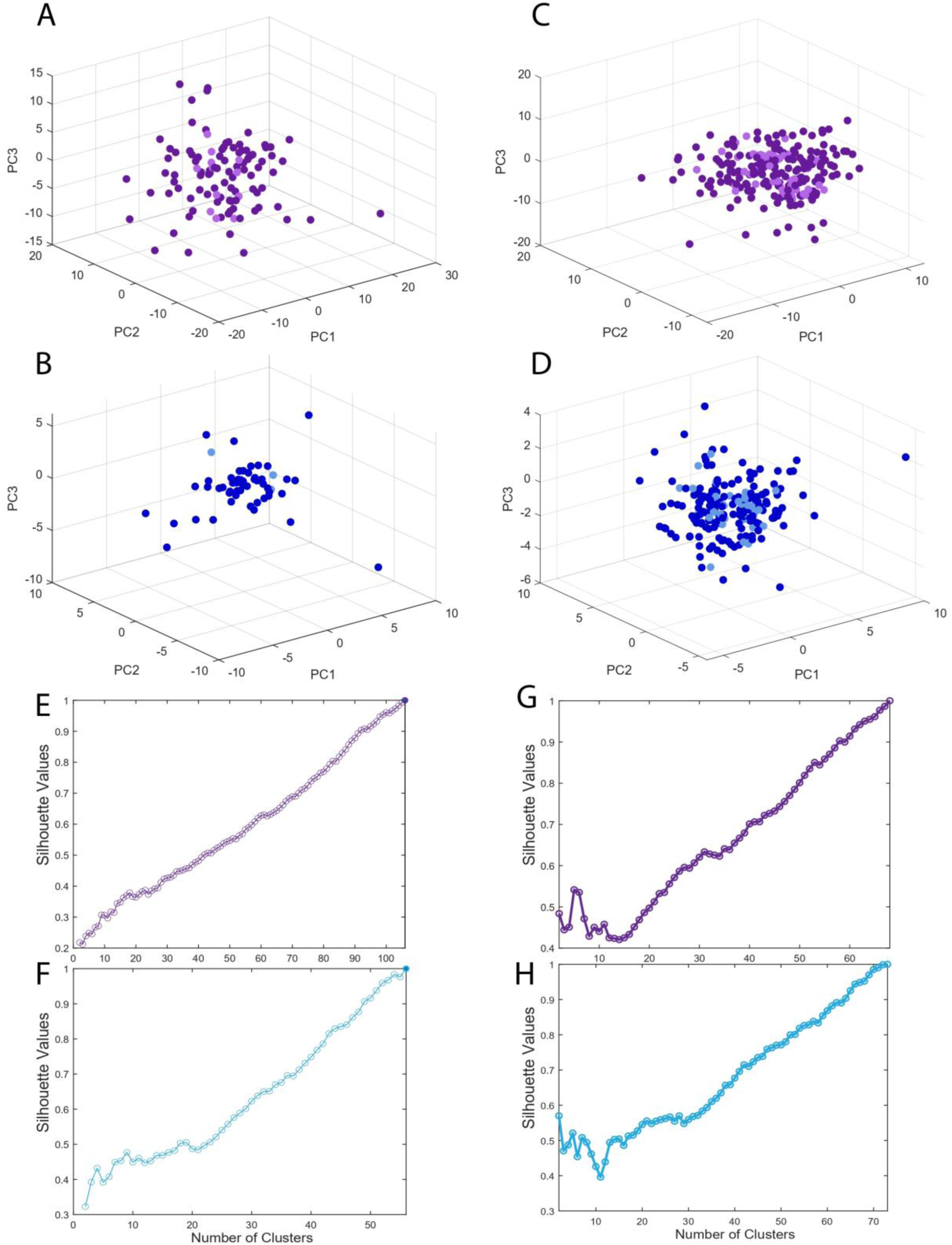
PCA and clustering analyses do not reveal discrete clusters of GP or M1 responses. (A–D) Principal component analyses (PCA) of normalized residual responses. Each dot represents a single unit, plotted along the first three principal components. Darker colors represent responses in the overlearned condition, and lighter colors represent responses in the random condition. (A) GP units from monkeys E and F. (B) M1 units from monkeys E and F. (B) GP units from monkeys C and H. (D) M1 units from monkeys C and H. (E–H) Silhouette analyses of clustering applied to normalized residual responses. Curves show silhouette values as a function of cluster number, with higher values indicating stronger clustering. (E) GP units from monkeys E and F, overlearned condition. (F) M1 units from monkeys E and F, overlearned condition. (G) GP units from monkeys C and H, overlearned condition. (H) M1 units from monkeys C and H, overlearned condition.

We next asked whether hierarchical clustering could uncover categorical response motifs.

Residual response profiles were clustered using cosine similarity, and the optimal number of clusters was estimated from silhouette values, which measure how well each unit matches its assigned cluster relative to other clusters, with higher values indicating more distinct cluster separation(Rousseeuw 1987). Under a start/stop framework, one would expect a small number of clusters, with large groups of neurons exhibiting peaks aligned to the first or last sequence element. Contrary to this prediction, the silhouette analysis produced no local maxima, indicating that no particular cluster number was favored. Instead, silhouette values increased linearly with each additional cluster (Figure 7E–H), implying that the data are better characterized as a continuum of overlapping response patterns rather than discrete categories. For monkeys C and H, the silhouette curve showed a modest elbow, reflecting a change in slope, but the values did not peak and decline. Instead, they continued to increase monotonically with additional clusters, indicating that no particular cluster solution was favored.

Together, these analyses argue against the idea that GPi or M1 neurons are organized into distinct start-and stop-related classes. Instead, both regions appear to represent sequence order through a distributed, heterogeneous code that tiles the entire sequence space.

## DISCUSSION

We set out to test whether neurons in the primate basal ganglia exhibit boundary-selective “bracketing” signals or whether they encode sequence order more uniformly across the sequence. Contrary to predictions from boundary-specific models, we found no evidence for preferential activity at sequence initiation or termination in either GP or M1. Instead, many neurons in both regions encoded ordinal position, with responses tiling the entire sequence space. This representation was more prominent in overlearned sequences but was also detectable under a random condition in which the sequence element order was unknown.

Prominent work in rodents has inspired models in which the basal ganglia initiate and terminate motor programs. Evidence for this model has come from studies that found that striatal projection neurons, dopaminergic neurons, and neurons in GPe displayed, to a varying degree, selective modulation at sequence boundaries. This selectivity developed as mice learned a lever-press sequence (Jin and Costa 2010, Jin, Tecuapetla et al. 2014) or navigated a T-maze (Barnes, Kubota et al. 2005, Thorn, Atallah et al. 2010). We reasoned that if start-and stop-signals are present in the striatum, the input nucleus of the BG, they should also be preserved in GPi, the output nucleus. We did not observe this. Rather, we observed a novel result: after regressing out information associated with kinematics, the residual spike counts of neurons in GP encoded the ordinal position of all sequence elements uniformly. Furthermore, this encoding was seen to a similar degree in M1 as well. Although encoding was far more prominent during the performance of fixed sequences, it was also present in a small number of neurons during the performance of a random sequence.

The representation of serial order by single-units has long been thought to be accomplished primarily by cells in premotor regions, such as the supplementary motor area.(Clower and Alexander 1998, Shima and Tanji 2000) Recent work in humans has reported a role for M1 in representing single elements of a fixed, button-press sequence (Yokoi, Arbuckle et al. 2018, Beukema, Diedrichsen et al. 2019). Prior work has also found neurons in primate GPi that are preferentially active at certain phases of a motor sequence (Mushiake and Strick 1995).

Our findings build on this work by showing that this representation is also found at the single unit level and is independent of kinematics.

Two classes of models may account for these results. In “BG-tutors-cortex” models, BG output provides teaching signals that help cortex acquire sequence representations (Turner and Desmurget 2010). Alternatively, in “cortex-tutors-BG” models, cortical inputs drive basal ganglia activity during sequence learning (Kawai, Markman et al. 2015) until skilled movement become cortex-independent. Our results cannot decisively distinguish between these possibilities, but the presence of order coding in both regions suggests an interactive process that results in shared representations of sequence-related information. The lack of discrete boundary signals argues against models in which GP output gates entire motor programs and instead supports models where BG output contributes to continuous monitoring and structuring of serial order.

The patterns in neural activity we observed parallel the animals’ behavioral performance. In the fixed condition, reaction times were markedly shorter and movements more stereotyped, consistent with chunking or habit formation. Neural tiling of sequence elements in both GP and M1 may provide the scaffolding for this smoother performance. This resonates with broader views of the basal ganglia as essential not for initiating actions but for shaping the efficient expression of well-practiced skills (Dhawale, Wolff et al. 2021).

Several limitations warrant consideration. Recent work has suggested a privileged role for the BG in performing sequences in the absence of visual cues (Mizes, Lindsey et al. 2023). One potential criticism of this study is that the performance of the overlearned sequence was always sensory guided, rather than internally generated. We do not believe this would significantly impact our interpretation of the behavior as a skill or habit. The performance of most skills is often accompanied by sensory feedback. This feedback allows for the erroneous performance of the skill to be halted. We believe that having a task that features visual cues for each movement to be ethologically sound. Furthermore, in the overlearned sequence, reaches often begin with a short enough latency relative to cue onset that it is physiologically impossible for the cue to be instrumental in aiding the initial generation of the movement. Ongoing studies in our lab will concretely address the difference in M1 and GP activity during the performance of overlearned sequences with and without visual cues.

We did not record enough trials to have each movement direction represented an adequate number of times at each ordinal position in the random sequence. This would have allowed for a more direct comparison of the two conditions when the movement direction and ordinal position in the sequence are the same. Also, we only commenced recording once the fixed sequence was overlearned. It is of interest to record activity while animals learn a fixed sequence to examine the timescale on which this ordinal position encoding develops and if it occurs first in cortex or in GP. Ongoing studies in our lab will address both issues.

In summary, our findings demonstrate that GP and M1 neurons encode ordinal position throughout motor sequences, with uniform responses tiling the sequence space. These results challenge models that cast the basal ganglia as mere initiators or terminators of motor programs and instead suggest a broader role in representing serial order alongside cortex. This distributed coding may be fundamental to how the brain handles the serial order problem in motor behavior.

## ACKNOWLEDGMENTS

We thank Lisa Nieman-Vento and Cherie Lee Cornmesser for their contributions to animal care. Research reported in this publication was supported by the National Institutes of Health under award numbers R01NS39146, P01NS044393, and R01NS113817. The content is solely the responsibility of the authors and does not necessarily represent the official views of the National Institutes of Health.

